# Identification of *Bulinus forskalii* as a potential intermediate host of *Schistosoma hæmatobium* in Senegal

**DOI:** 10.1101/2022.06.15.496218

**Authors:** Papa Mouhamadou Gaye, Souleymane Doucouré, Doudou Sow, Cheikh Sokhna, Stéphane Ranque

## Abstract

Understanding the transmission of *Schistosoma hæmatobium* in the Senegal River Delta requires knowledge of the snails serving as intermediate hosts. Accurate identification of both the snails and the infecting *Schistosoma* species is therefore essential. Cercarial emission tests and multi-locus (COX1 and ITS) genetic analysis were performed on *Bulinus forskalii* snails to confirm their susceptibility to *S. hæmatobium* infection. A total of 55 *B. forskalii*, adequately identified by MALDI-TOF mass spectrometry, were assessed. Cercarial shedding and RT-PCR assays detected13 (23.6%) and 17 (31.0%), respectively, *B. forskalii* snails parasitised by *S. hæmatobium* complex fluke. Nucleotide sequence analysis identified 6 (11.0%), using COX1, and 3 (5.5%), using ITS2, *S. hæmatobium*, and 3 (5.5%) *S. bovis*. This result is the first report of infection of *B. forskalii* by *S. hæmatobium* complex parasites.

## Introduction

Snails of the genus *Bulinus* (Müller, 1781) are intermediate hosts for the larval development of trematodes parasite species of the *Schistosoma hæmatobium* species complex in Africa, the eastern Mediterranean, and Madagascar [1]. Several species of the S. *hæmatobium* group are endemic in Africa, notably *S. hæmatobium, S. intercalatum, S. guineensis, S. bovis* and *S. curassoni*. Only the first three are involved in human diseases [2,3]. Urogenital schistosomiasis, caused by *S. hæmatobium* (Bilharz, 1852), is present in most countries on the African continent [4], especially in all regions of Senegal [5]. *S. hæmatobium* is highly endemic in sub-Saharan Africa, where 600 million people are at risk of infection, and 200 million urinary schistosomiasis cases [6], including 150,000 deaths [7], are recorded *per* year.

In endemic areas, transmission of *S. hæmatobium* involves various freshwater gastropod snails species of the genus *Bulinus* [8–10]. In Senegal, *Bulinus truncatus, B. senegalensis, B. globosus* and *B. umbilicatus* are the main intermediate hosts of *S. hæmatobium* [9,11,12]. *B. forskalii* (Ehrenberg, 1831) is also present in temporary pools and, especially in the Senegal River, in *S. hæmatobium* permanent transmission areas [10,13]. *B. forskalii* is often misidentified as *B. senegalensis*, a species with which it shares similar morphological features and often the same biotopes [14]. These two species are genetically similar and form, with *B. camerunensis*, the *B. forskalii* group [14,15]. The presence of *Bulinus forskalii* has been reported to be involved in *S. bovis, S. guineensis*, and *S. intercalatum* biological cycle in Western and Central Africa [3,16,17]. Whereas this Bulinid species has never been reported to be infected with *S. hæmatobium* in Senegal [18], previous studies have indicated that *B. forskalii* could be a potential intermediate host for *S. hæmatobium* in Niger [19]. The main limitation of these results was that snails identification relied on morphological features only, and thus identification errors cannot be excluded due to the known morphological similarity between *B. forskalii* and *B. senegalensis* [15].

## Methods

In September 2020, we collected (n=55) snails from the Senegal River Delta area, identified as *B. forskalii* (**Fig.1**) using the Mandahl-Barth identification key based on shell morphology [20]. We then performed cercarial shedding tests. The excreted cercariae were not kept. Then, each snail specimen was dissected for Desoxyribo Nucleic Acid (DNA) extraction and Matrix Assisted Laser Desorption/Ionization-Time of Flight (MALDI-TOF MS) analysis. The feet of the snails were used for MALDI-TOF MS identification as described by Hamlili *et al*. (2021) [21]. Each specimens’ MALDI-TOF MS spectra were then compared to the in-house snails reference spectra database (available at https://doi.org/10.35088/f605-3922)[21] using the MALDI-Biotyper v3.0 software (Bruker Daltonics). The blind test identification quality is estimated via log score values (LSV), which quantifies the degree of identity between the query and the reference spectra in the database. A sample is considered correctly identified when the LSV value is ≥ 1.7. The remaining body of each specimen was used for genomic DNA extraction. The *Bulinus forskalii* MS identification was confirmed by nucleotide sequence-based analysis of the Folmer region of cytochrome C oxidase (COX1) subunit I, using the universal primers LCO1490, and HCO2198 [22,23]. Real-time-Polymerase Chain Reaction (RT-PCR) targeting the *S. hæmatobium* group specific Dra1 repeat region sequence[24,25] was performed on *B. forskalii* DNA extracts to assess infection with *S. hæmatobium* group parasites. PCR sequencing of COX1 [26] and the second internal transcribed spacer (ITS2) of the rRNA gene [27] was performed to identify the species within the *S. hæmatobium* group.

**Fig 1.**
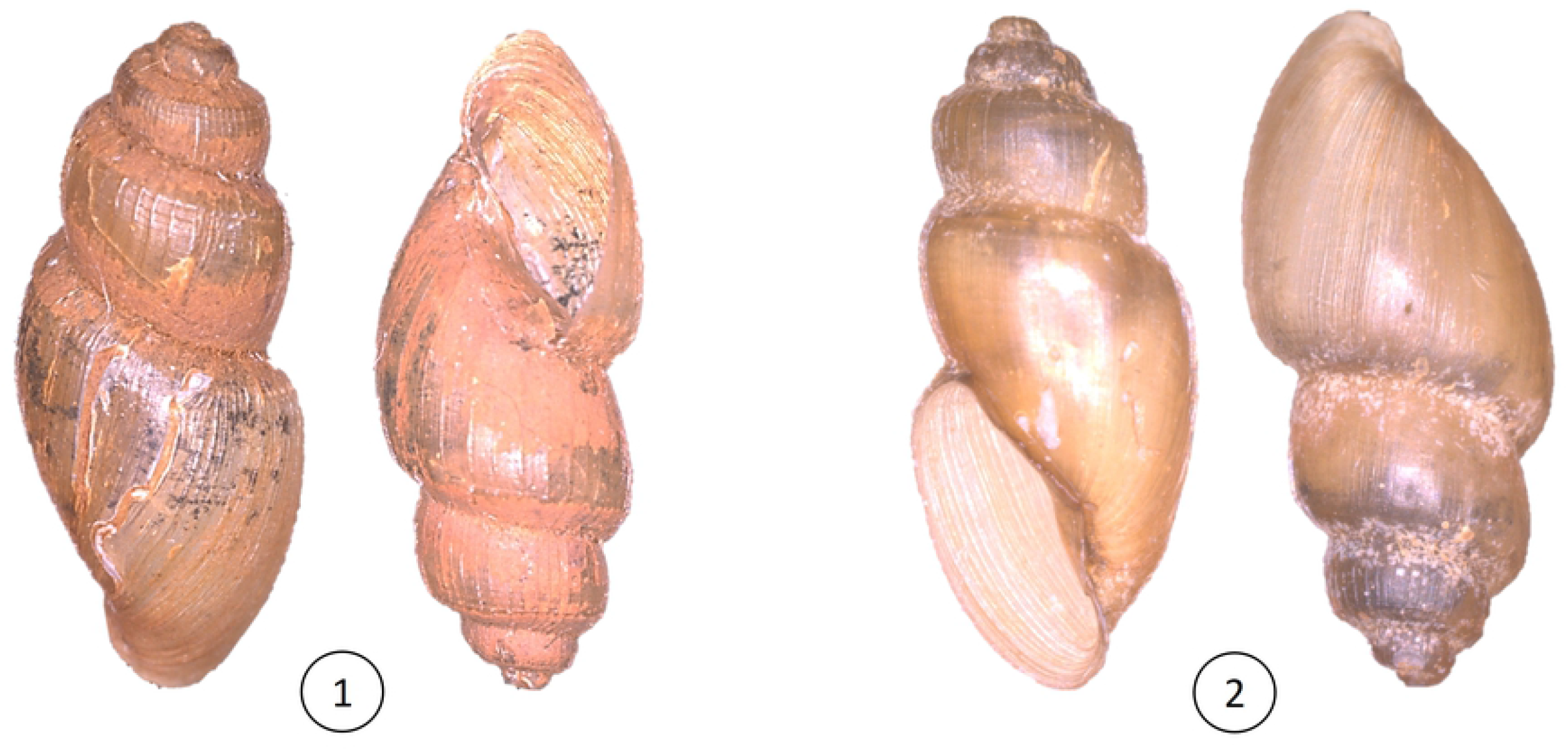
Morphological features of the *Bulinus forskalii* snail species analysed in the current study.

## Results

All snails were identified as *Bulinus forskalii* by MALDI-TOF MS with log scores ranging from 1.89 to 2.63 [mean (SD) = 2.31 (0.20)] and reproducible spectra, indicative of a reliable identification (**Fig.2**). *Bulinus forskalii* identification were further confirmed by 100.0% identity with the COX-Folmer of *Bulinus forskalii* (GenBank Accession No: M535893.1) [3,22]. In the 55 *B. forskalii* specimens, 13 (23.6%) and 17 (30.9%) were infected with *Schistosoma* parasites with the cercarial shedding tests or the Dra1-RT-PCR, respectively. The RT-PCR was positive (Cycle threshold (Ct) ≤ 35) in each snail that shed cercariae (**Table 1**). The COX1 nucleotide sequence further identified 6 (10.9%) *S. hæmatobium*, and 3 (5.5%) *S. bovis*. In contrast, ITS2 nucleotide sequence-based identification yielded 3 (5.5%) for both *S. hæmatobium* and *S. bovis*. Basic Local Alignment Search Tool (BLAST) query of the National Center for Biotechnology Information (NCBI) nucleotide database identified either *S. hæmatobium* or *S. bovis*, as detailed in Table 1.

**Fig 2.**
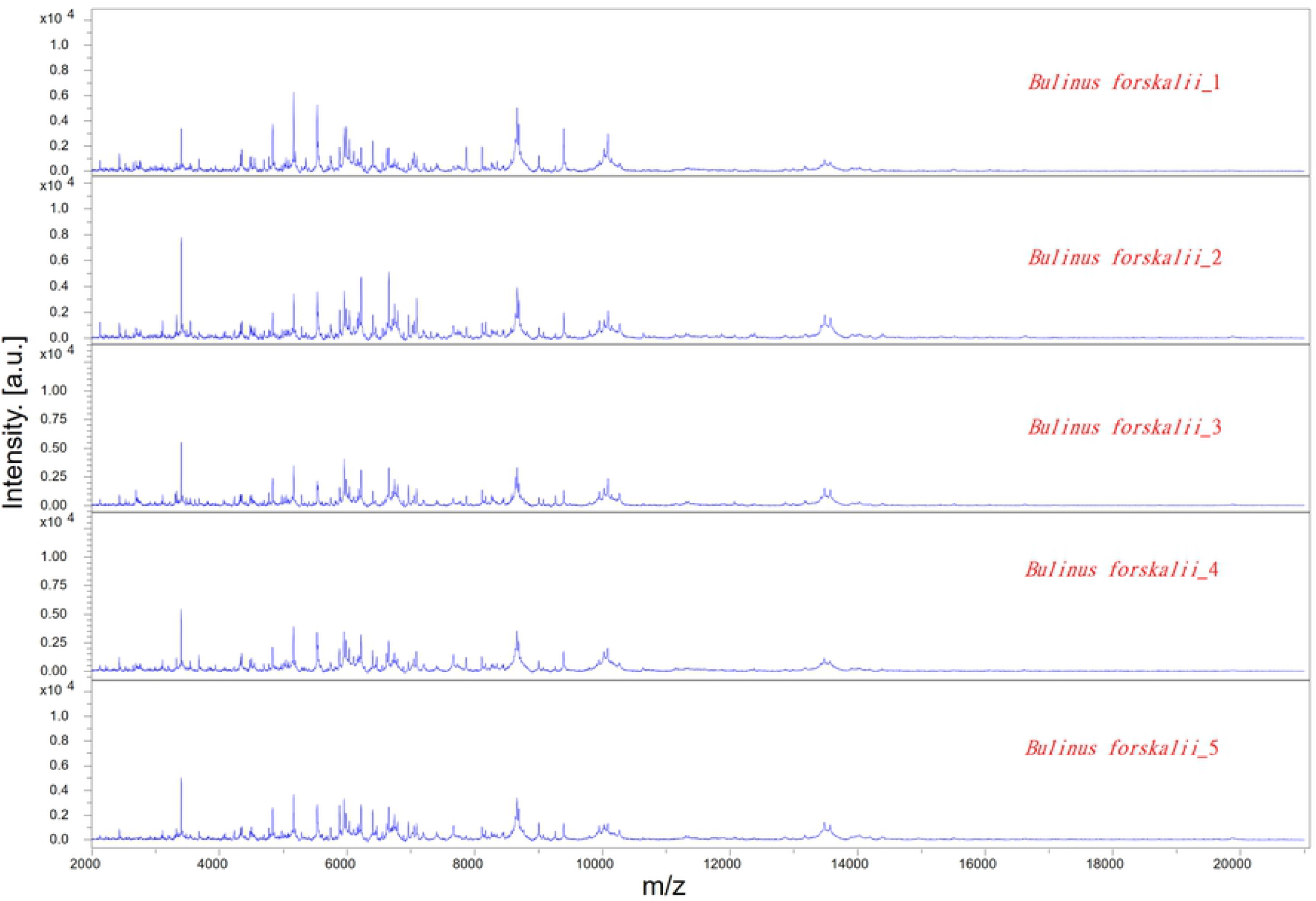
Characteristic MALDI-TOF MS spectra acquired from *Bulinus forskalii* feet.

**Table 1.** MALDI-TOF MS identification of the snail parasitised by *S. hæmatobium* complex fluke, and COX1 and ITS2 DNA sequence-based identification of the *Schistosoma* spp. parasites.

| Samples ID | Snail species ID by MS | LSV | cercarial shedding test* | RT-PCR (Ct) | Schistosomes species ID by Cox1 (% Identity) | GenBank Accession No | Schistosomes species ID by ITS2 (% identity) | Accession |
| --- | --- | --- | --- | --- | --- | --- | --- | --- |
| KABfI4 | *Bulinus forskalii* | 2.618 | (+) | 29.43 | *S. hæmatobium* (100.00%) | MT579447.1 | NA** | - |
| KABfI5 | *Bulinus forskalii* | 2.367 | (+) | 29.23 | *S. hæmatobium* (99.78%) | MT579447.1 | NA | - |
| KABfNI5 | *Bulinus forskalii* | 1.889 | (-) | 28.57 | *S. hæmatobium* (99.78%) | MT579447.1 | NA | - |
| S20BfI7 | *Bulinus forskalii* | 2.553 | (+) | 23.01 | *S. hæmatobium* (99.78%) | MT579447.1 | *S. bovis* (100.00%) | MT580958.1 |
| S20BfI9 | *Bulinus forskalii* | 2.564 | (+) | 31.17 | *S. hæmatobium* (99.78%) | MT579447.1 | NA | - |
| S20BfI10 | *Bulinus forskalii* | 2.625 | (+) | 25.00 | NA | - | *S. hæmatobium* (100.00%) | MT580959.1 |
| KSD2BfI1 | *Bulinus forskalii* | 2.288 | (+) | 19.39 | *S. bovis* (99.23%) | MT159594.1 | *S. bovis* (100.00%) | MT580958.1 |
| KSD2BfI4 | *Bulinus forskalii* | 2.377 | (+) | 25.41 | *S. bovis* (99.61%) | MT159594.1 | *S. bovis* (100.00%) | MT580958.1 |
| S9BfNI1 | *Bulinus forskalii* | 2.313 | (-) | 34.94 | *S. hæmatobium*<br>(100.00%) | MT579447.1 | *S. hæmatobium*<br>(100.00%) | MT580959.1 |
| SOBfNI1 | *Bulinus forskalii* | 2.107 | (-) | 33.88 | *S. bovis* (99.61%) | MT159594.1 | *S. hæmatobium*<br>(100.00%) | MT580959.1 |

\*Cercarial shedding test: (+), positive; (-), negative.
\*\* NA: not available, due to sequencing failure

## Discussion and Conclusion

Studies aiming to detect and identify parasites of the *S. hæmatobium* group in snails are scarce in Senegal. Our findings demonstrate that *B. forskalii* can be parasitised by *S. hæmatobium* complex fluke, in particular, *S. hæmatobium* s.s., *S. bovis* s.s., and hybrids between them. Our findings contrast with those of Tian-Bi *et al*. [28] who found no infected *B. forskalii* after the cercarial shedding test. Nevertheless, studies conducted in *B. senegalensis* and *B. umbilicatus* in central Senegal have shown similar results [11]. The evidence of *S. hæmatobium* cercariae excretion by *B. forskalii* demonstrates that this snail acts as intermediate host, enabling the evolution from the miracidium to the cercaria stages of the parasite.

The evidence that *B. forskalii* can be an intermediate host of *S. hæmatobium* complex, parasites that are involved in schistosomiasis in Senegal, is a significant contribution to improving our knowledge on host-parasite interactions that should be taken into account in further epidemiological studies and schistosomiasis control programs. The main limitation of our study is that discrepant ITS2 and COX1-based identification in a given snail specimen can be interpreted either as an infection with one hybrid parasite or a co-infection with two distinct parasite species. Further studies on isolated cercariae should give a final answer this open question.

